# Uncovering the Design Rules for Sustainable Growth of Mineralized Mycomaterials

**DOI:** 10.1101/2025.09.12.675881

**Authors:** Dylan H. Moss, Olivia Pear, Jorge Guío, Alyssa Libonati, Daniel Ducat, Konane Bay, Arjun Khakhar

## Abstract

Mycomaterials, materials made from filamentous fungi, have several advantages over traditional materials, such as their genetic programmability and self-healing properties. However, their lack of mechanical strength and cost of production often constrain the applications they can be used in. In this work, we take inspiration from natural systems to overcome these challenges by elucidating design principles for mineralization-based enhancement of mechanical strength and synthetic lichen-based low-cost growth. We demonstrate that surface display of an enzyme from sea sponges, silicatein α, on the hyphae of the filamentous fungus *Aspergillus niger* enables mineralization of polysilicate and that this does not impact fungal growth. We also show that this strategy can be extended to other silicatein α variants and characterize how the degree of mineralization can be modulated. We then demonstrate that mineralization enhances the mechanical properties of the mycelium, including its tensile strength, modulus, and toughness. Finally, we show how these reinforced mycelia can be grown without external carbon sources using a synthetic lichen-based co-culture to facilitate low cost biomanufacturing. Together, our results lay the groundwork for the sustainable production of mineralized mycomaterials and create a new model system to study how mineralization impacts growth and mechanical properties.

**Significance Statement:** Materials made from filamentous fungi, called mycomaterials, have several advantages over traditional materials, but their poor mechanical properties and relatively high production costs have limited their application. We elucidated design principles to enable tunable mineralization of fungal mycelium and have shown that it does not impact growth but significantly enhances mechanical strength. We have also shown that these reinforced mycelia can be grown without any external carbon in a synthetic lichen like consortia to minimize production costs. This work creates a novel experimental system to study how mineralization impacts growth and mechanical properties and will facilitate the broader application of mycomaterials in the future.

## Introduction

Materials built from the mycelium of filamentous fungi, termed mycomaterials, present several advantages over traditional materials including programmable microstructure, self-assembly, and self-healing(1). However, their limited mechanical strength and toughness have constrained their application to soft foams or malleable leathers(2). In advanced polymeric materials high mechanical strength and toughness are often achieved through embedding inorganic components within an organic matrix to create a nanocomposite. The hard inorganic constituents reinforce the strength and toughness of the softer organic matrix(3, 4). Prior studies have sought to generate such composites by growing mycelium through porous rigid substrates and then baking them. While this strategy does improve the material’s strength(5), it is resource intensive and thus not easily scaled, which has limited its broad application. It is also challenging to maintain consistent properties and to predictably modulate them due to the amorphous and non-homogeneous nature of substrates(6).

An alternative strategy to generate mechanically strong mycomaterials which does not suffer these limitations is to genetically encode the capacity to form organic-inorganic composites into the fungi. There are many examples across the clades of life where such composites are generated through enzymatic mineralization to provide strength, spanning human bones, starfish ossicles, and sea sponge exoskeletons(7–9). In the case of sea sponges an abundant mineral in sea water, silica, is mineralized into amorphous silica oxide, termed bioglass, by a class of enzymes called silicateins(10). Silicatein α, one of several silicateins that are embedded in the protein filaments used by sea sponges to produce bioglass, has been previously characterized as sufficient to catalyze the mineralization of ionic silica *in vitro* using a synthetic substrate called tetraethyl orthosilicate (TEOS)(11) . The active site of this enzyme contains a catalytic triad of residues which facilitate the hydrolysis of TEOS via a nucleophilic attack on the silicon atom, releasing ethanol. The hydrolyzed silanol side chain is then able to undergo polycondensation via dehydration with other free silanols, forming polysilicate(12, 13).

Pioneering work in bacteria and yeasts has shown that display of these enzymes on the surface of cells results in deposition of amorphous polysilicate on the cell surface(14, 15). While this has been leveraged to create lenses and facilitate aggregation, thus far these approaches have not been explored in the context of generating organic-inorganic composites to enhance mechanical properties.

In addition to suboptimal material properties, the cost associated with growing mycomaterials is another barrier to their broad application. Driving fungal growth necessitates an external carbon source, which is generally a central cost to biomanufacturing microbe-based materials(16). However, natural systems have evolved methods to generate utilizable carbon sources from the abundant resources of carbon dioxide, water, and sunlight, namely photosynthesis. Prior work has shown how a photosynthetic microbe, *Synechococcus elongatus* PCC 7942, can be engineered to share its fixed carbon to drive the growth of co-cultured heterotrophs(17).

In this work, we elucidate a set of design principles to genetically encode the capacity to form polysilicate coatings on the hyphal surface in the model filamentous fungus *Aspergillus niger*. We first explore how the formation of this coating impacts fungal growth. We then go on to characterize how silicatein α expression, substrate concentration, and exposure time impact the degree of mineralization in these strains. We also test whether this strategy is modular enough to be extended to other silicatein α variants.

Finally, we characterize how this mineralization alters the mechanical properties of the mycelium and demonstrate how such materials can be grown sustainably using a synthetic lichen-based co-culture with engineered cyanobacterium.

## Results

### Surface display of silicatein α enables mineralization of fungal hyphae

We first sought to explore if it was possible to genetically encode polysilicate deposition on the cell surface via surface display of silicatein α (SilA) on the hyphae of the model filamentous fungus *Aspergillus niger.* This fungal chassis was chosen due to its demonstrated capacity for high levels of protein secretion, thereby enabling significant surface expression of SilA. We designed a chimeric SilA fusion protein consisting of a secretion tag(18), the SilA enzyme from *Latrunculia oparinae* (LoSilA)(19), and a cell wall binding protein (20) separated by flexible linkers (**Fig. 1A**). This gene was assembled into an expression cassette driven by the promoter of the glucoamylase gene, which shows strong and carbon source dependent expression(21, 22) thereby enabling easily tunable expression, using Golden Gate assembly. It was subsequently loaded into a fungal entry vector that had a constitutive PyrG expression cassette to enable auxotrophic selection.

**Figure 1.**
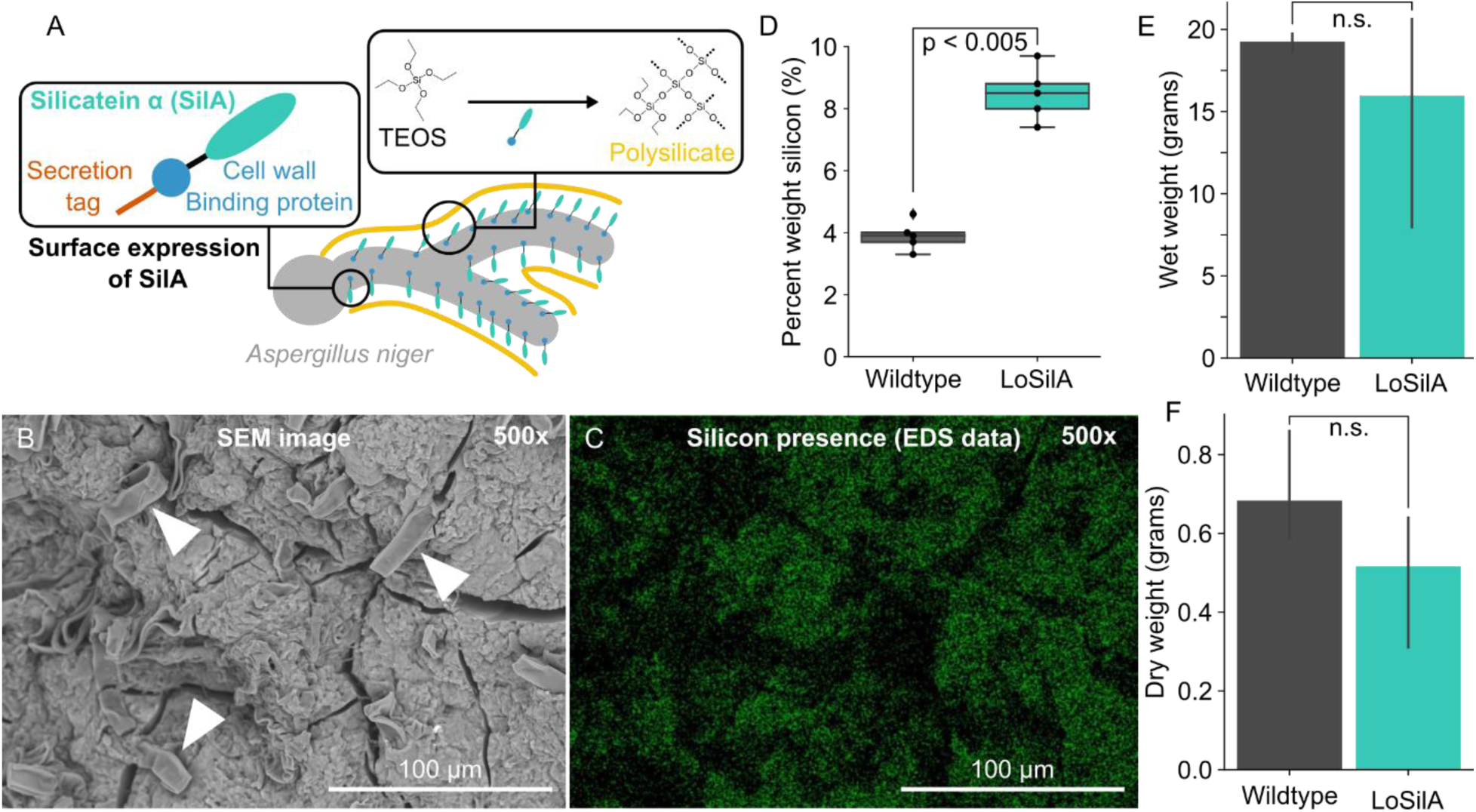
Surface display of Silicatein α in *A. niger* enables mineralization of polysilicate without impacting growth. A) Schematic depicting the fusion protein consisting of a secretion tag, cell wall binding protein, and silicatein was expressed in *A. niger* to enable surface display of Silicatein α (SilA) (left box). This leads to TEOS condensation into polysilicate (right box). B) Representative SEM image of a SilA expressing strain treated with 50mM TEOS for 5 days showing fungal hyphae (white arrows) embedded in mineralized polysilicate. C) Paired EDS data for the SEM image in B where the green dots indicate the presence of a silicon signal. D) Box plot summarizing EDS measurements of average percent weight of silicon deposited on the surface of either Wildtpe (Gray) or LoSilA expressing (teal) fungi. Each dot represents a single biological replicate. E,F) Bar plots summarizing measurements of wet (E) and dry (F) weight of either wildtype (gray) or LoSilA expressing (teal) fungi grown for seven days in CM media with 50mM TEOS. All plots constitute three biological replicates and n.s. stands for not significantly different.

This vector was linearized and transformed into *A. niger* protoplasts using established protocols(23). Positive transformants were subsequently assayed for their capacity to enable mineralization compared to the parental untransformed strain.

To test mineralization, we grew up mycelium of these strains from spores in submerged liquid culture under sufficient shaking to generate spherical pellets. After 2 days of growth in complete media (CM), we moved them to sterile deionized water containing 50mM TEOS for one week under the same growth conditions. These samples were subsequently dried and the percentage weight of silicon deposited on the hyphal surface was characterized by scanning electron microscopy (SEM) and energy dispersive X-ray spectroscopy (EDS)(15). Imaging revealed clear evidence of polysilicate deposition on the hyphal surface of LoSilA expressing strains that were treated with TEOS (**Fig. 1B,C**). We observed that in areas where hyphae were densely packed it appeared that the hyphae were embedded within a polysilicate matrix. However, we did not observe similar deposition on fungi not expressing SilA that was treated with TEOS or on the LoSilA untreated samples, demonstrating this effect is driven by the surface expressed SilA (**Fig S1**). As secondary confirmation, our EDS data shows approximately 2 fold higher percent weight of silicon on mycelium expressing LoSilA compared to wildtype controls treated with TEOS (**Fig 1D**). Taken together these results demonstrate that surface expression of SilA is sufficient to generate a mineralized mycelium.

### Hyphal mineralization does not compromise growth

We next tested if mineralization impacted mycelium growth, for example by forming a rigid casing that prevents hyphal extension. We inoculated flasks of CM with spores from the SilA expressing strain or the wildtype control and grew them for 2 days, after which we added TEOS to a final concentration of 50mM and grew them for 7 more days.

Subsequently the growth was quantified by measuring both the wet and dry weight of the mycelium. We observed that the SilA expressing strain had wet and dry weights that were not significantly different from the control (**Fig 1E,F**), which demonstrates that mineralization of polysilicate on the hyphal surface does not impede mycelial growth.

### The degree of mineralization can be modulated by altering the level of SilA expression using alternative carbon sources

We hypothesized that as the polysilicate mineralization was an enzymatically catalyzed process, altering the expression of the SilA should enable modulation of the mycelial mineralization. Prior work has shown that the glucoamylase promoter we used to drive LoSilA expression is induced by maltose leading to increased levels of protein production(21, 22). We leveraged this inducible system to modulate LoSilA expression and test its impact on mineralization (**Fig 2A**). We grew fungal cultures in CM that had glucose replaced with an equal amount of maltose for 2 days and then transferred them to a solution of deionized water with 50mM TEOS for 7 days. These cultures were then dried and the percentage weight of silicon deposited on their surface was analyzed with EDS. We observed a significant increase in silicon in the SilA expressing strains grown in maltose as compared to glucose (**Fig 2B**). This demonstrates that the degree of mycelial mineralization generated by our system can be tuned through modulation of SilA expression via changes to the media carbon source.

**Figure 2.**
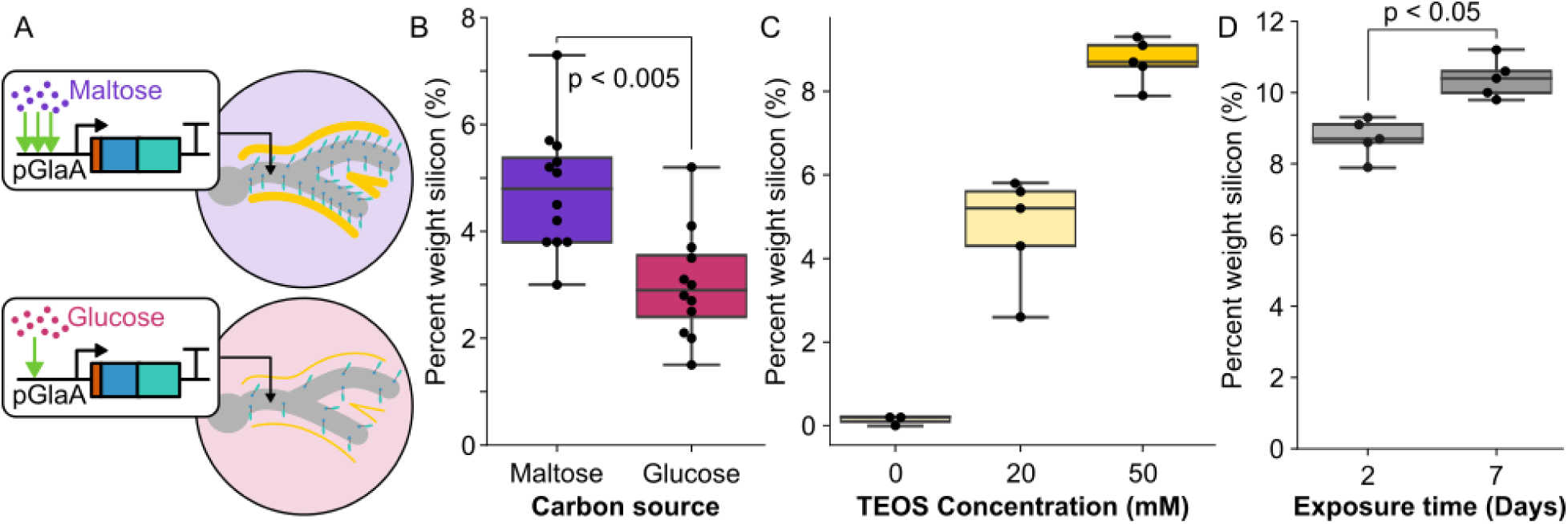
The degree of mycelial mineralization can be tuned. A) Graphic depicting that the GlaA promoter (pGlaA) used to drive the SilA fusion protein is more strongly induced by maltose (purple) than glucose (pink). B) Boxplots summarizing EDS measurements of the percent weight of silicon on the surface of the LoSilA expressing strain grown in CM media supplemented with either maltose (purple) or glucose (pink) and then treated with 50mM TEOS for 2 days. C) Boxplots summarizing EDS measurements of the percent weight of silicon on the surface of the LoSilA expressing strain grown in CM media supplemented with maltose and then treated with either 0, 20, or 50mM TEOS for 2 days. D) Boxplots summarizing EDS measurements of the percent weight of silicon on the surface of the LoSilA expressing strain grown in CM media supplemented with maltose and then treated with 50mM TEOS for either 2 or 7 days. Each dot represents an independent biological replicate.

### The degree of mineralization depends on substrate concentration and exposure time

We next set about characterizing how the substrate concentration and exposure time to the substrate impacted the degree of mycelial mineralization. To do this, we grew cultures of the SilA expressing strain and a wildtype control in tandem in CM with either glucose or maltose as the carbon source for 2 days and then transferred them to DI water with either 0mM, 20mM or 50mM TEOS and incubated for either 2 or 7 days. Individual mycelial pellets were then extracted, dried, and analyzed with EDS. For samples grown in maltose we observed significant increase in mineralization with both increasing substrate concentration (**Fig 2C**) and longer exposure time at 50mM TEOS (**Fig 2D**). While we observed self-condensation of TEOS on the wildtype control with high concentrations of TEOS and long exposure times, consistent with previous mineralization studies that use this substrate, the increase in mineralization on the SilA expressing strain was consistently greater (**Fig S2**). Similar trends, with lower effect sizes, were observed in glucose, which is consistent with what we would expect for strains with lower expression of SilA (**Fig S2**).

Interestingly, in the samples grown in maltose and treated with 20mM TEOS, we observed a significant decrease in mineralization between two and seven days (**Fig S2**). To further understand the dynamics of mineralization at a 20mM substrate concentration, we repeated this experiment with a higher resolution time course where mineralization of the mycelium was measured daily. We observed that peak mineralization occurred at 2 days of exposure to TEOS, after which there was a steady decline in the degree of mineralization with day 5 being significantly lower than day 2 and only marginally above background levels (**Fig S3**). We hypothesized that this effect might be due to the acidification of media by *A. niger,* which is known to secrete significant quantities of organic acid, as a lower pH would initially disfavor the precipitation of TEOS into polysilicate and potentially etch away existing polysilicate from the surface of hyphae. Additionally, prior studies have shown that *A. niger* can solubilize insoluble silicate from quartz(24). When we measured pH of the media used for mineralization, we observed significantly lower pH for later timepoints, which is consistent with this hypothesis (**Fig S3**). We hypothesize that this effect is not visible in the 50mM TEOS treatments as the higher rate of mineralization associated with increased substrate concentration counteracts the impact of acidification. Taken together, these results demonstrate that both substrate concentration and exposure time can be modulated to control the degree of mineralization.

### Surface displayed SilA enzymes can be modularly swapped

Prior studies have characterized a range of SilA variants from different sea sponges with different catalytic rates and substrate specificities(11, 25–27). If the SilA displayed in our system was modular, it could be swapped to enable the construction of diverse organic-inorganic composites in the future. To test the modularity of the system, we built a strain that expressed a modified version of the chimeric fusion protein with LoSilA replaced with the SilA from *Tethya aurantium* (TaSilA) using the techniques described previously (**Fig 3A**). The mineralization of this strain was then compared to previously tested strains, the LoSilA expressing strain and our control, by growing them in tandem in maltose containing CM media and then transferring them to DI water with either 20mM or 50mM TEOS for two days. These samples were subsequently dried and the degree of mineralization was quantified via SEM-EDS as described previously. In both conditions we observed that the TaSilA expressing strain had a significantly higher percentage weight of silicon compared to the wildtype control demonstrating that mineralization was occurring (**Fig 3B-G**, **S1,S4**). We also observed that the degree of mineralization was not significantly different from that of LoSilA in either condition (**Fig 3B,C**). While the absolute degree of mineralization LoSilA in these experiments was lower than previous experiments, likely due to differences in TEOS stocks used and fungal growth prior to mineralization, the fold change in comparison to the wildtype control grown in tandem was consistent with previous observations lending confidence to these conclusions. These results demonstrate that the surface expressed SilA can be easily swapped to alter catalytic rate and substrate specificity in the future.

**Figure 3.**
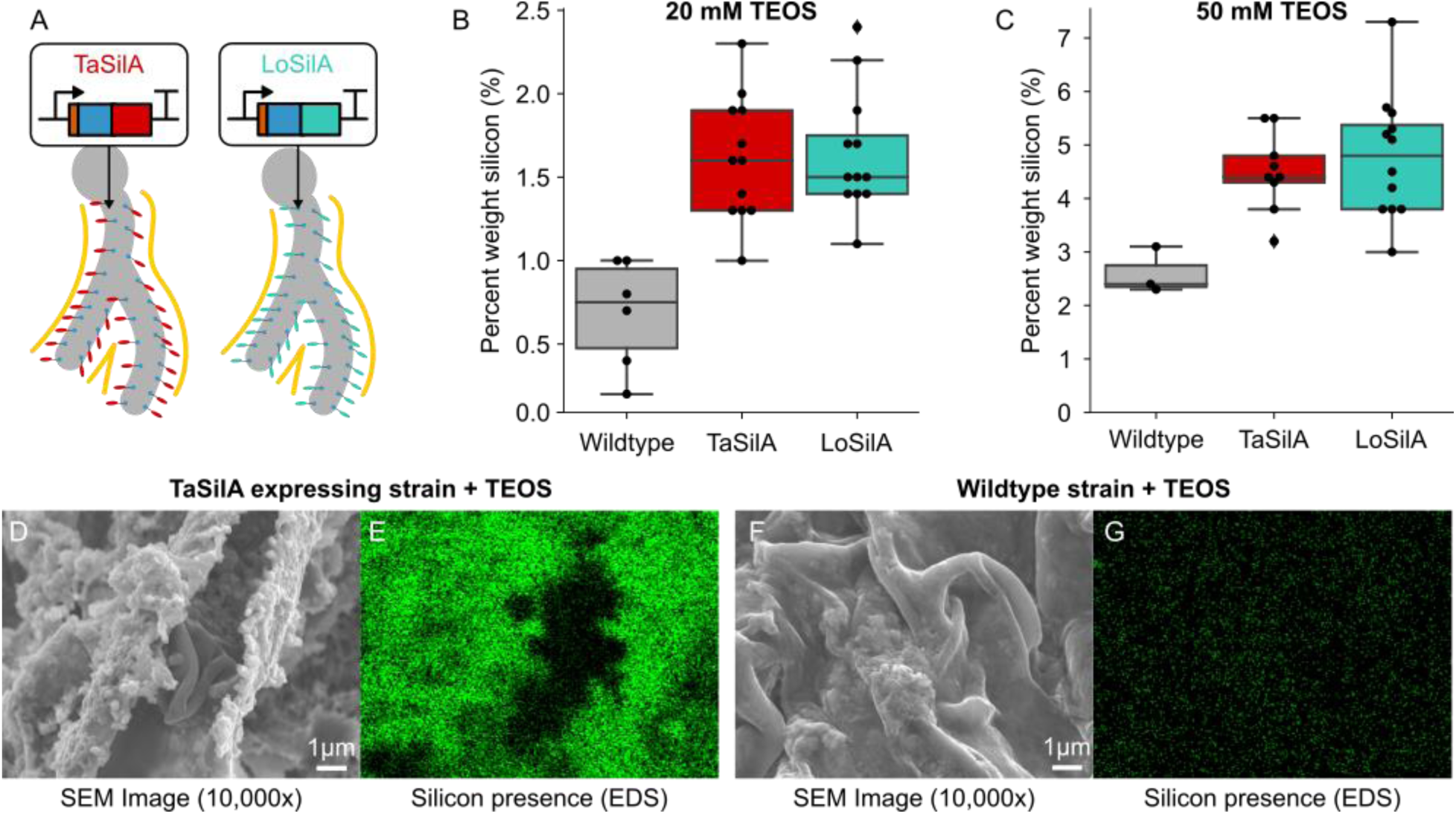
The SilA being displayed on the hyphal surface can be modularly swapped. A) Graphic depicting how the LoSilA in the chimeric fusion protein can be modularly swapped with TaSilA. B,C) Boxplots summarizing EDS measurements of the percent weight of silicon on the surface of wildtype (gray), TaSilA expressing fungi (red), or LoSilA expressing fungi (teal) grown in CM media supplemented with maltose and then treated with either 20mM (B) or 50mM (C) TEOS for 2 days. Each dot is an independent biological replicate. D) Representative SEM image of a TaSilA expressing strain treated with TEOS showing fungal hyphae embedded in mineralized polysilicate. E) Paired EDS data for the SEM image in D where the green dots indicate the presence of a silicon signal. F) Representative SEM image of a wildtype strain treated with TEOS showing a lack of mineralized polysilicate. G) Paired EDS data for the SEM image in F where the green dots indicate the presence of a silicon signal.

### Mineralization of mycelium enhances strength

Organisms in nature utilize mineralization as a strategy for developing strength and toughness from soft building blocks. The production of a mineral fraction from organic components, such as those of nacre, coral, and bone, leads to improved mechanical properties of the resultant composite structure(28, 29). Based on our understanding of these naturally mineralized materials, we hypothesize that the production of silica deposits by the fungus will result in increased mechanical strength at the macroscale. Having demonstrated that *A. niger* can be engineered to generate tunable organic-inorganic composites via polysilicate mineralization, we next set about characterizing how this mineralization impacted the mycelium’s mechanical properties at the macroscale using uniaxial tensile tests. Spores of either LoSilA expressing strains or a wildtype control were grown in CM containing glucose for six days to generate pellets approximately 25mm in diameter. These were then mineralized via two days of exposure to 20mM hydrolyzed TEOS and then pressed into films. A set of non-mineralized controls were pressed immediately after growth. All samples were conditioned overnight to 55% relative humidity. These samples were then cut into 20 mm x 5 mm rectangular strips, and their tensile properties were tested using a TA.XTPlusC texture analyzer.

The mineralized LoSilA expressing strain films demonstrate statistically significant increases in Young’s modulus (*E*), ultimate tensile strength (UTS), and toughness compared to the non-mineralized LoSilA films (**Fig 4A-D**). The increase in mechanical properties due to mineralization is observed in the representative stress-strain curves for LoSilA expressing strain films with and without TEOS exposure (**Fig 4A**). The mycelium films demonstrate non-linear stress-strain behavior, with curves containing a linear elastic region at low strain and a strain hardening region at intermediate strains until failure (**Fig 4A**). The mineral volume fraction contributes to the stiffness of the mycelium according to the rule of mixtures(30, 31). We used the rule of mixtures, or Voigt model(32), to approximate the volume fraction of silica in the mineralized LoSilA expressing mycelium film as 8.8 % (LoSilA film exposed to 20 mM TEOS for 48 hours). We utilized thermogravimetric analysis (TGA) to assess the Voigt model prediction, and the data show good agreement of 8.4% residual mass (**Fig S5**). Among the fungal strains tested, the non-mineralized samples do not demonstrate statistically significant differences in *E* or UTS, suggesting that the surface display of LoSilA in the absence of its substrate, TEOS, does not alter the mechanical properties significantly (**Fig S5**). The *E* and UTS of wildtype *A. niger* films exposed to TEOS also do not exhibit significant differences compared to non-mineralized films, suggesting that spontaneous precipitation of TEOS in the absence of SilA is not sufficient to alter the mycelium’s mechanical properties (**Fig S5**). These data together suggest that the mineralization of polysilicate on the LoSilA mycelium films enhances their mechanical strength and stiffness.

**Figure 4.**
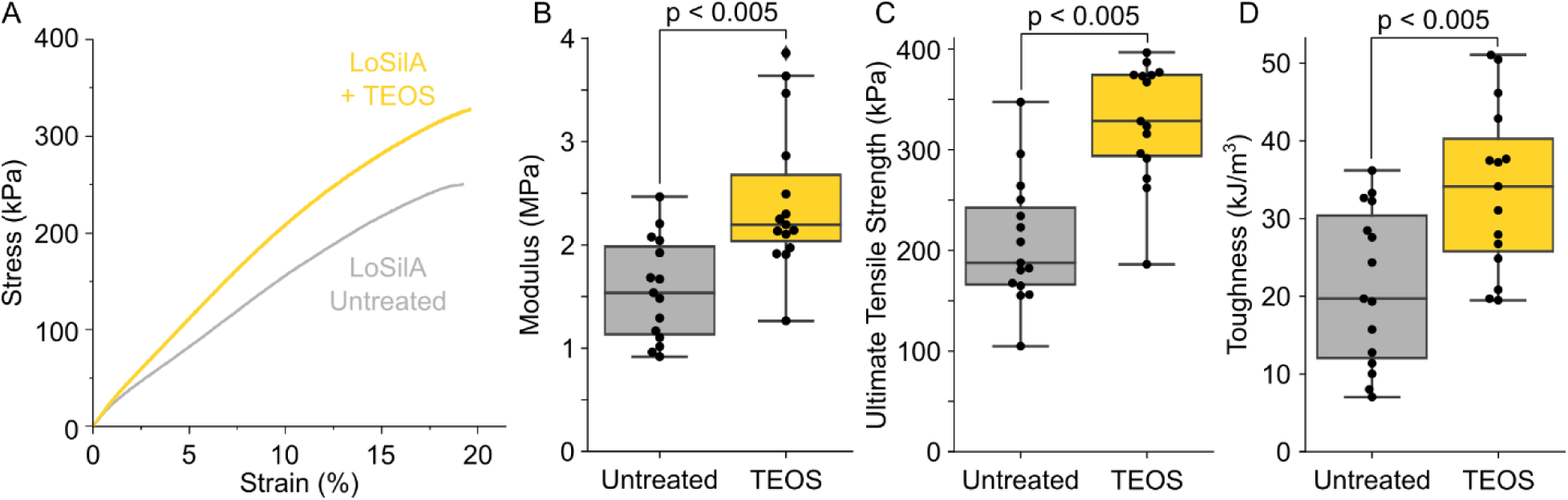
Mineralization of mycelium enhances its mechanical strength. A) A representative stress-strain curve of a LoSilA expressing strain that is either treated with 20mM TEOS for two days (yellow) or that is untreated (gray). B-D) Box plots summarizing measurements of Young’s modulus (B), Ultimate tensile strength (C), and Toughness (D) calculated from stress-strain curves of a LoSilA expressing strain that is either treated with TEOS (yellow) or that is untreated (gray). Each dot represents an independent biological replicate.

### Mineralizing mycelium can be grown autotrophically via co-culturing with sucrose secreting cyanobacteria

The mineralization-based enhancement of mechanical properties we have engineered makes mycomaterials a promising alternative to conventional materials. However, their production must also be cost-competitive for their application to be viable. Since carbon inputs necessary to drive microbial growth are a major contributor to production costs(33), we hypothesize that minimizing these inputs would enable sustainable and cost-effective production of mineralized mycomaterials. To achieve this, we aimed to establish a synthetic lichen-like co-culture comprising our *A. niger* LoSilA strain together with cyanobacterium *Synechococcus elongatus* PCC 7942 *cscB/sps* strain.

This *cscB/sps* strain was previously engineered to secrete a portion of its photosynthetically fixed carbon as sucrose via IPTG based induction of a *sucrose permease* (*cscB*) and a *sucrose phosphate synthase* (*sps*) genes (17) (**Fig 5A**). We hypothesized that this co-culture would enable fungal growth without the need for external carbon supplementation, thereby enabling sustainable biomanufacturing of mineralized mycomaterials.

**Figure 5.**
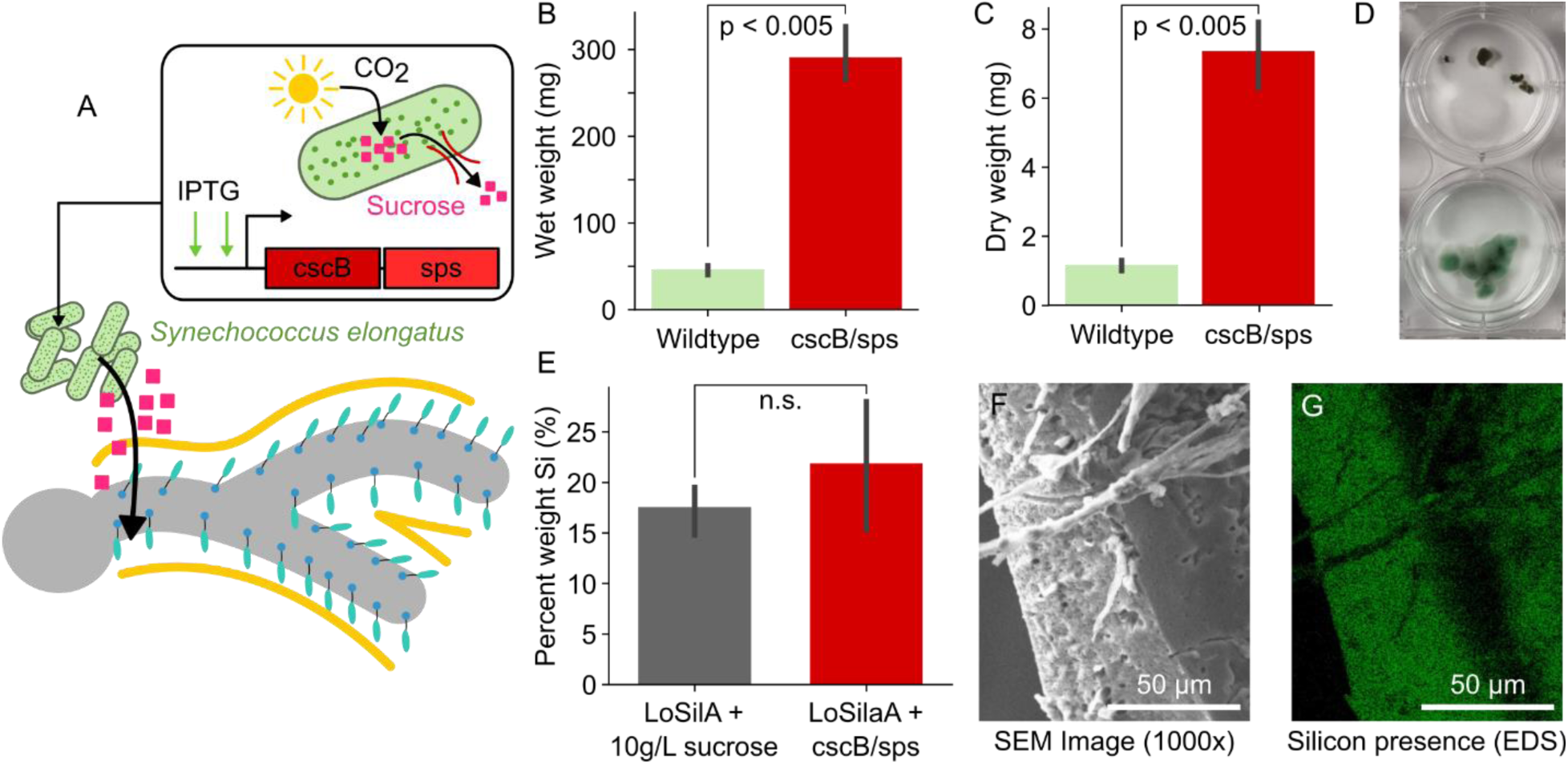
Synthetic lichen-based co-cultures enable autotrophic growth of mineralized mycelium. A) Schematic of engineered *S. elongatus* strains expressing *cscB* and *sps* (red) in response to IPTG induction. This leads to secretion of a fraction of their photosynthate as sucrose into the surrounding media, thereby creating a carbon source to drive growth of mineralizing mycelium. B,C) Bar plots summarizing measurements (n=3) of wet (B) and dry (C) biomass of LoSilA expressing *A. niger* co-cultured with either wildtype (gray) or *cscB/sps* (red) *S. elongatus*. D) Representative images of LoSilA expressing fungi co-cultured with either wildtype (top) or *cscB/sps* (bottom) *S. elongatus* prior to measurement. D) Bar plots summarizing EDS measurements (n=3) of the percent weight of silicon on the surface of LoSilA expressing *A. niger* grown with either sucrose as a carbon source (gray) or in co-culture with *cscB/sps S. elongatus* (red) and treated with 50mM TEOS for 5 days. “n.s.” indicates no statistically significant difference. F) Representative SEM image of LoSilA expressing fungi in co-culture with *cscB/sps S. elongatus* showing hyphae embedded in a polysilicate matrix. G) Paired EDS data for the SEM image in F where the green dots indicate the presence of a silicon signal

We first quantified sucrose secretion by the engineered *cscB/sps* strain in BG11 co-culture medium (BG11[CO]) after 3 days of IPTG induction. Sucrose levels reached 0.88 ± 0.007 g/L, whereas no sucrose was detected in wild-type (WT) controls under the same conditions. We then established co-cultures by combining either whole cyanobacterial cultures or cell-free supernatants from *cscB/sps* cultures induced for 3 days with LoSilA spores germinated in MM for 3 days. Fungal growth was determined after 10 days by measuring wet and dry biomass.

Both co-cultures with *cscB/sps* cells (**Fig 5 B, C, D**) and *cscB/sps* cell-free supernatants (**Fig S6 B, D, F**) supported LoSilA fungal growth, with significantly higher fungal biomass than co-cultures with WT *S. elongatus* cells or cultures with cell-free supernatants of WT *S. elongatus*. In fact, while fungal growth in co-cultures with the WT strain was similar to growth in BG-11[CO] without any supplemental carbohydrate source (**Fig S6 A, C, E**), fungal growth in co-cultures with the *cscB/sps* strain reached 6-7 fold higher total biomass: nearly half the growth observed in BG-11[CO] supplemented with 10 g/L sucrose (**Fig S6, A, C, E**). These results indicate that sucrose secreted by *cscB/sps* strain was sufficient to sustain fungal growth.

Importantly, in addition to supporting fungal growth, we also observed strong mineralization in co-cultures of LoSilA with *cscB/sps* strain without any external carbon source, with silicon levels comparable to those of cultures in BG11[CO] with 10 g/L sucrose (**Fig 5 E, F, G, S7**). This demonstrates that incorporating cyanobacteria into the mycelium to enable lichen-like growth does not hinder SilA-driven polysilicate mineralization. Taken together, these results demonstrate that photosynthetically derived sucrose from engineered *cscB/sps S. elongatus* can serve as the sole carbon source for sustainable production of mineralized mycomaterials.

## Discussion

Diverse natural systems utilize mineralization to generate organic-inorganic hybrid materials with the strength to support their development and to provide resilience to external forces. Recapitulating this behavior synthetically in fungal mycelium would expand the applications of mycomaterials to situations that require mechanical strength. Our results demonstrate that the display of SilA on the hyphal surface enables polysilicate mineralization without impacting fungal growth. This is in contrast to previous studies that have surface-displayed SilA on single celled microbes, where mineralization created a rigid barrier that inhibited cell division thereby halting growth(15). We hypothesize that, as the SilA is fused to the secretion signal from the GlaA gene, it is likely secreted mostly from the growing hyphal tip like the native gene(34). Once secreted we expect the cell wall binding domain also fused to SilA directs binding to the hyphal surface distal to the tip, where mineralization subsequently takes place. This would enable the growing hyphal tips to extend while the mycelium becomes mineralized facilitating tandem growth and mineralization.

Our results show that there are several different avenues to tune the degree of mineralization including changing the substrate concentration, the exposure time, or the expression of SilA. This highlights how the growth and mineralization conditions of a single strain could be modulated to achieve a range of bulk material properties. The dependence of mineralization on SilA expression, which we modulated via changes to the media’s carbon source, demonstrates that differential mineralization across the mycelium could be genetically encoded in the future to achieve more complex material properties. Our results also showed that *A. niger*’*s* propensity to acidify its growth media inhibits mineralization at lower substrate concentrations. Prior studies have described how specific genetic knockouts can reduce organic acid secretion(35), which could mitigate this issue in the future.

We also demonstrate that our system is modular, enabling the SilA displayed on the hyphal surface to be easily swapped with other variants. As prior studies have described a range of naturally evolved and engineered SilA enzymes with different catalytic rates(25), this provides another avenue to tune the degree of mineralization. This also opens the door to mineralizing a range of other metal oxides, including those of titanium, zirconium, gallium, barium, cerium, and calcium, as SilA variants have been shown to accept a range of different substrates(25–27, 36). Thus, the design principles we elucidate could be applied in future studies to generate more complex organic-inorganic hybrid materials with novel material properties.

By demonstrating that the mineralization of mycelium significantly enhances the tensile properties and strength of mycelial films, this work lays the groundwork for expanding mycomaterials to applications where strength is necessary. We expect that the size and morphology of mineral deposits would alter how they impact the mechanical properties of the film. More in-depth characterization of these relationships in future studies would facilitate better forward engineering of mycomaterials. Additionally, prior studies have shown that polysilicate mineralization can have several other interesting material properties beyond mechanical reinforcement such as being good insulators and focusing light(15, 37). These could be the basis of exciting future applications of these mineralized mycomaterials.

Our work showing that these mineralized mycomaterials can be grown in co-culture with an engineered cyanobacterium as its sole carbon source highlights the potential for sustainable biomanufacturing of these materials. While we use a relatively expensive SilA substrate, TEOS, for experimental convenience, sea water is an abundant source of ionic silicon and the substrate for SilA in its native context. Prior studies have shown that both *A. niger* and *S. elongatus* can be engineered to be tolerant to sea water(38, 39) to enable low-cost mineralization. However, this would need to be experimentally validated in future studies. The combination of lichen-like autotrophic growth of mycelium coupled with the mechanical enhancement conferred by mineralization could enable leveraging this system for generating self-standing 3D biological structures in the future, which has thus far remained a grand challenge in the field of engineered living materials. However, while these experiments serve as a proof-of-concept, characterization of the impact of mineralization on cyanobacterial photosynthesis and the stability of these co-cultures over several months is necessary to determine if they could be applied to generate large scale structures in the future. Additionally, while we chose to use *A. niger* due to its highly efficient protein secretion, we hope to explore if this system could be extended to other fungi used in mycomaterials, such as *Ganoderma lucidum* and *Trichoderma reesei,* in the future.

Taken together, our results elucidate the design principles for mineralized mycomaterials that can be sustainably manufactured and demonstrate that this mineralization enhances their tensile strength. In addition to its translational value, our work creates an exciting new model system for future studies into how mineralization impacts 3D growth and how the interplay of morphology and mineralization impacts structural properties at the macro scale.

## Materials and Methods

### Construction of *Aspergillus niger* strains with surface expressed silicatein α

All work was done with strain FGSC A1179 (MA70.15) procured from the Fungal Genetics Stock Center (FGSC) at Kansas State University.(40) A1179 has the following genotype (ΔkusA, pyrG-, (cspA1)). (41) This strain is a derivative of strain N402. Plasmids encoding expression cassettes for SilA surface expression were built using Golden Gate Assembly using a combination of synthesized sequences and parts from the Modular Synthetic Biology Toolkit for Filamentous Fungi.(42) To build this we first built a fungal entry vector that used a pCambia backbone with homology regions that are 500bp upstream and downstream of the native pyrG locus to enable HDR mediated knock-in.(43) This vector included the *Aspergillus fumigatus*’s pyrG coding sequence and terminator upstream of the 5’ homology region to enable pyrG based selection.

We then constructed a level 1 expression vector that encoded an expression cassette for the SilA chimeric fusion protein driven by the *A. niger* glucoamylase (glaA) promoter (pglaA) (21, 22). We domesticated this promoter by removing a PaqCI site which would interfere with our assembly process. The chimeric fusion protein consists of the *A. niger* glaA secretion signal sequence(18), the silicatein α enzyme from TaSilA or LoSilA(19) codon optimized for *A. niger* using Integrated DNA Technologies’ Codon Optimization Tool, a 5x GS linker, a 6x HisTag, 2x GGGS linker, and the glycosylphosphatidylinositol-anchored cell wall mannoprotein (cwpA) from *A. niger*(20). This is followed by the *Penicillium rubens* OAT1 terminator from the kit and then the 500bp downstream homology region. Full sequences can be found in **Table 1**. A level 2 assembly was used to construct the final transformation template. This was verified via whole plasmid sequencing prior to transformation.

**Table 1.**
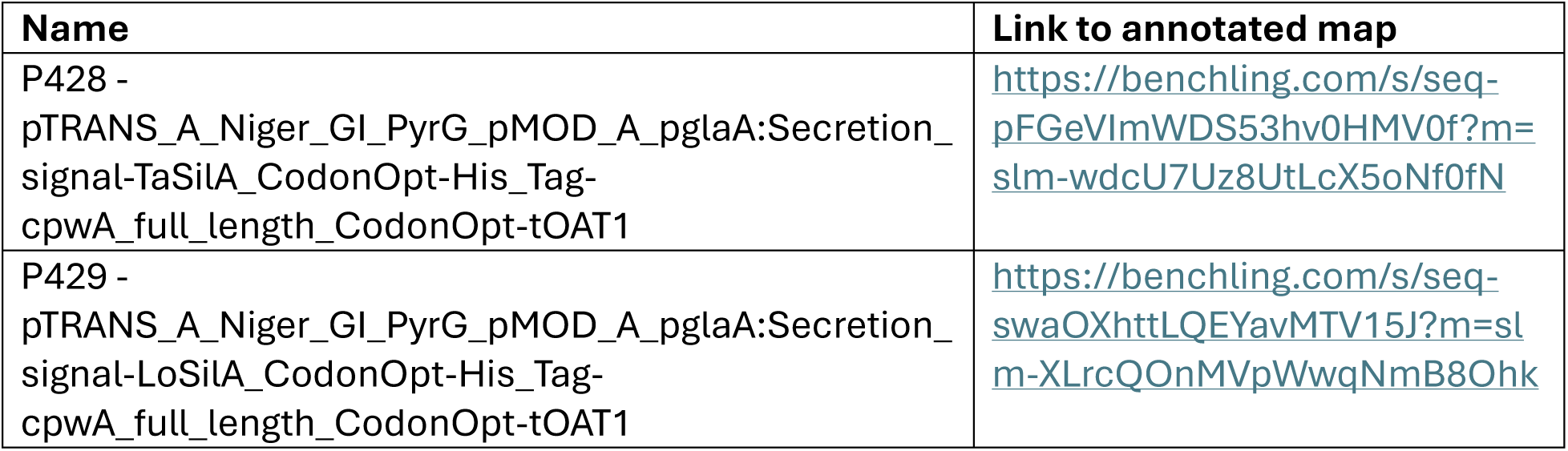
List of plasmids used in this study.

### Transformations

Final constructs containing the entire insert region were then amplified with Q5® High-Fidelity DNA Polymerase from the beginning of the 500bp upstream to the end of the 500bp downstream homology region. A portion of the PCR product was run on a gel to confirm that fragments were the correct size, and then the linearized DNA fragments were used for transformations. Protoplast mediated transformations were performed using standard protocols with linear DNA fragments(23). Primary transformants were passaged onto Minimal Media. Transformants were then passaged to 5mL liquid CM without uracil and uridine and grown for 2 days in culture tubes at 37C and 150RPM. DNA was then extracted using previously published protocols(44) . PCR verification was then performed using Q5® High-Fidelity DNA Polymerase.

### Fungal Growth Conditions

Cultures were grown in Complete Media and Minimal Media for Aspergillus as described. (45) Complete and Minimal Media were supplemented with the described Vitamin Supplement. For transformations and screening Complete Media was prepared as described. For experimental work comparing strain performance, Complete Media was supplemented with 5mM uracil and uridine (UU) to support the growth of untransformed A1179 (KO/WT) as our control. For experiments altering carbon source, maltose was used in the place of glucose (10g/L of glucose (CGM) or maltose (CMM)). For incubation in TEOS, sterile water (autoclaved MilliQ filtered H2O) was used. TEOS, a commonly used synthetic substrate for biomineralization studies, was slowly added to the H2O. Standard growth conditions for cultures entailed growth from spores in 50mL of CM+UU in 125mL Erlenmeyer flasks, 150RPM for 3 days. For experiments, CM was modified from the described recipe which uses 10g/L of glucose (CGM) to replace it with 10g/L of maltose (CMM).

### Biomass Measurements

Cultures were grown in 50mL CGM for 2 days at 150RPM and 30C. TEOS was then added in to the CGM for a concentration of 50mM. Cultures grew for 7 more days. Cultures were then poured out over Miracloth and transferred to pre-weighed weigh boats. Wet weight was then recorded. Weigh boats were then covered in tinfoil and transferred to an oven (70C) and dried for one week. Samples were then removed from the oven, tin foil was removed, and cultures were weighed. All p values to determine statistical significance were calculated using a student’s t-test. All data analysis and plotting were done in python and the relevant code can be found at (https://github.com/arjunkhakhar/250905_Mineralized_mycomaterials).

### Scanning Electron Microscopy (SEM) and Energy Dispersive X-ray Spectroscopy (EDS/EDX)

For further silication experiments, cultures were started from spores and grown in CGM+UU or CMM+UU for 2 days at 30C and 150RPM. These cultures were then poured out over Miracloth and the mycelial biomass was then transferred to a second set of flasks with autoclaved MilliQ water. TEOS was then added at specified concentrations (20mM or 50mM). Cultures were then incubated at 30C and 150RPM for the specified amount of time.

For SEM EDS analysis, individual pellets of fungus were extracted from the cultures and placed on double-sided carbon tape (PELCO Tabs™, Carbon Conductive Tabs, 15mm OD) on 15×10mm aluminum SEM mounts (933414 Sigma-Aldrich). These samples were then enclosed in a petri dish with micropore tape and transferred to an oven and dried for 2 days at 70C. Samples were coated with a 4nm layer of platinum with a Cressington Sputter Coater 108 Auto. Samples were imaged and analyzed with Hitachi TM4000Plus II. Higher magnification images were taken with JEOL 6500F and Hitachi SU3500. Accelerating voltage of 15keV was used for EDS analysis on the TM4000Plus II. Overview scans of the pellets to examine the amount of Si on the surface of the mycelium were performed at 25x magnification, and the outlines of each pellet were traced in Oxford Instruments’ AZtecOne EDS analysis software. This analysis tool returns the % Weight of detected atoms on the surface of a sample and were examined for Silicon (% Wt Si). The % Wt Si was then compared across strains and treatments. All data analysis and plotting were done in python and the relevant code can be found at (https://github.com/arjunkhakhar/250905_Mineralized_mycomaterials). All p values to determine statistical significance were calculated using a student’s t-test.

### pH Testing

Cultures were added to H2O with 20mM of TEOS. 5mL of the supernatant of the culture was then taken and pH was measured with a pH meter. Measurements were taken from samples after the initial addition of the fungus, at 2 days, and then at 7 days. All data analysis and plotting were done in python and the relevant code can be found at (https://github.com/arjunkhakhar/250905_Mineralized_mycomaterials).

### Mechanical properties of mycelia films

*A. niger spores (*∼10^4^ spores) were inoculated into 15mL of complete media in 50mL falcon tubes. The tubes were incubated at 30 °C and 150 rpm on a tilted tube rack attached to a shaker plate for six days. After the initial growth, spheres were incubated for 48 hours at room temperature with 15 mL of 20 mM hydrolyzed TEOS in water adjusted to approximately pH 7 in 50 mL falcon tubes. Films were prepared from biomass cultures by pressing an individual mycelia sphere (approx. diameter 25 mm) between two Teflon coated glass slides with 0.1 mm cover slips placed either side of the sphere and held together with binder clips. All samples were conditioned overnight in a 55% relative humidity chamber, using a saturated solution of magnesium nitrate.

Samples were removed from the chamber and cut into 20 mm x 5 mm rectangular strips using a sharp blade. The thickness of each strip was determined with digital calipers, taking an average of three measurements across the length of the strip. Samples were mounted with tape onto paper frames for ease of attachment into tensile grips. With sample ends tightened in the lower and upper grips, paper frame edges were cut, and the tensile test initiated. A 5-kg load cell TA.XT*Plus*C texture analyzer with tensile grips was used to measure tensile properties of films with tests conducted at a constant speed of 5 mm/min. Force-displacement data generated by the instrument were converted to stress-strain using Matlab code. Young’s modulus, *E,* was determined from the linear region of the stress-strain curve, between 0.5 – 2.5 % strain. The ultimate tensile strength (UTS) was determined from the maximum stress of the specimen. The toughness was determined as the area under the stress-strain curve up to the UTS. All data analysis and plotting were done in python and the relevant code can be found at (https://github.com/arjunkhakhar/250905_Mineralized_mycomaterials). All p values to determine statistical significance were calculated using a student’s t-test.

### Thermogravimetric Analysis (TGA)

TGA was performed using a TA Discovery TGA 5500 to study the thermal decomposition of the inorganic-organic composite mycelium films. Samples of 5 – 10 mg, obtained after mechanical testing, were placed in platinum pans and heated from 25 to 800 ℃ with a ramp rate of 20 ℃/min. The tests were conducted under continuous flow of N2 gas at 20 mL/min. The mass is recorded as a function of temperature and the resultant curves aid in identifying decomposition temperatures of the organic mycelium. The corresponding residual mass was obtained at 800 ℃, after the combustion of organic materials is completed.

### Co-culture experiments

Co-culture experiments were performed with wild-type *Synechococcus elongatus* PCC 7942 and *S. elongatus* sucrose secreting strain *cscB/sps*(17). The *cscB/sps* strain harbors the sucrose permease gene (*cscB*) from *E. coli* under control of isopropyl β-d-1-thiogalactopyranoside (IPTG) inducible promoter P_trc_ integrated at Neutral Site 3 and the sucrose phosphate synthase gene (*sps*) from *Synechocystis* sp. PCC 6803 under control of P_trc_ promoter integrated at Neutral Site 2. Cyanobacteria strains and co-cultures were grown in BG-11 co-culture medium (BG-11[CO]), a modified BG-11(46) medium containing two-fold of NaNO_3_ (3 g/L), five-fold of KH_2_PO_4_ (0.15 g/L), five-fold of MgSO_4_ (0.375 g/L), five-fold of micronutrients and 3 g/L HEPES pH 8.3.

For co-culture establishment, *Aspergillus niger* LoSilA spores were germinated in 50 mL of Minimal Media (MM) at an initial density of 1·10^5^ spores/mL. Fungal cultures were maintained in 125 mL non-baffled Erlenmeyer flasks for 3 days at 30 °C and 100 rpm in an Innova 4230 incubator. Spores used to start cultures were collected from complete media (CM) agar plates using sterile water. In parallel, sucrose secretion was induced for 3 days adding 1 mM of IPTG to 50 mL cultures of *cscB/sps* in BG-11[CO] with initial OD_750_ of 0.3. 1mM was also added to WT *S. elongatus* PCC 7942 cultures, which served as control.

Cyanobacteria cultures were maintained in baffled 125 mL Erlenmeyer flasks for 3 days at 32 °C, 150 μmol photons m^−2^ s^−1^ of continuous light and 150 rpm in a Multitron incubator supplemented with 2% CO_2_. Prior to IPTG induction, cyanobacteria cultures were back-diluted daily to an OD_750_ of 0.3 for at least 3 days. Secreted sucrose was quantified from supernatants using the Sucrose/D-Glucose Assay Kit (K-SUCGL; Megazyme).

To establish co-cultures, MM was removed from *A.niger* germinated spores after 3 days and either whole cultures or supernatants of IPTG-induced WT and *cscB/sps S. elongatus* were poured into 125 mL non-baffled flasks containing the germinated spores. Co-cultures were maintained at 30 °C, 50 μmol photons m−2 s−1 of continuous light and 100 rpm in an Innova 4230 incubator for 10 days. For fungal growth determinations, wet and dry biomass were quantified as described previously. For silication assays, mycelial biomass was transferred to flasks with 50 mL of autoclaved MilliQ water and was treated with 20 mM of TEOS for 2 days or 50 mM of TEOS for 5 days. Silication was quantified as described previously.

## Acknowledgments

Jorge Guio and Daniel Ducat thank Greg Bonito and Bryan Rennick (MSU Department of Plant, Soil, and Microbial Sciences) for providing access to their facilities for fungal cultivation. Arjun Khakhar and Dylan Moss thank Ken Kassenbrock (CSU Department of Biology) for use of his biosafety cabinet. SEM/EDS was performed both at CSU ARC (RRID: SCR_021758) and we would like to thank Rebecca Miller for her assistance, as well as at COSINC-CHR CU Boulder (RRID:SCR_018985) and we would like to thank Adrian Gestos for his assistance. This work was supported by funds from the National Science Foundation grant 2334680 and 2516056 to Arjun Khakhar, grant 2516055 to Konane Bay, and grant 2334681 to Daniel Ducat.

## Author Contributions

DM built the strains, designed and executed all the experimental work associated with figures 1-3, and helped write the paper. AL designed and developed the experimental method associated with figure 4. OP executed all the experimental work associated with figure 4 and helped write the paper. JG designed and executed all the experimental work associated with figure 5 and helped write the paper. KB acquired funding to support the work, advised on experimental work associated with figure 4 and helped edit the paper. DD advised on experimental work associated with figure 5, acquired funding to support the work, and helped edit the paper. AK acquired funding to support the work, conceptualized the project, advised on the experimental work in figures 1-3, and wrote and edited the paper.

---

**Supplementary Figure S1.**
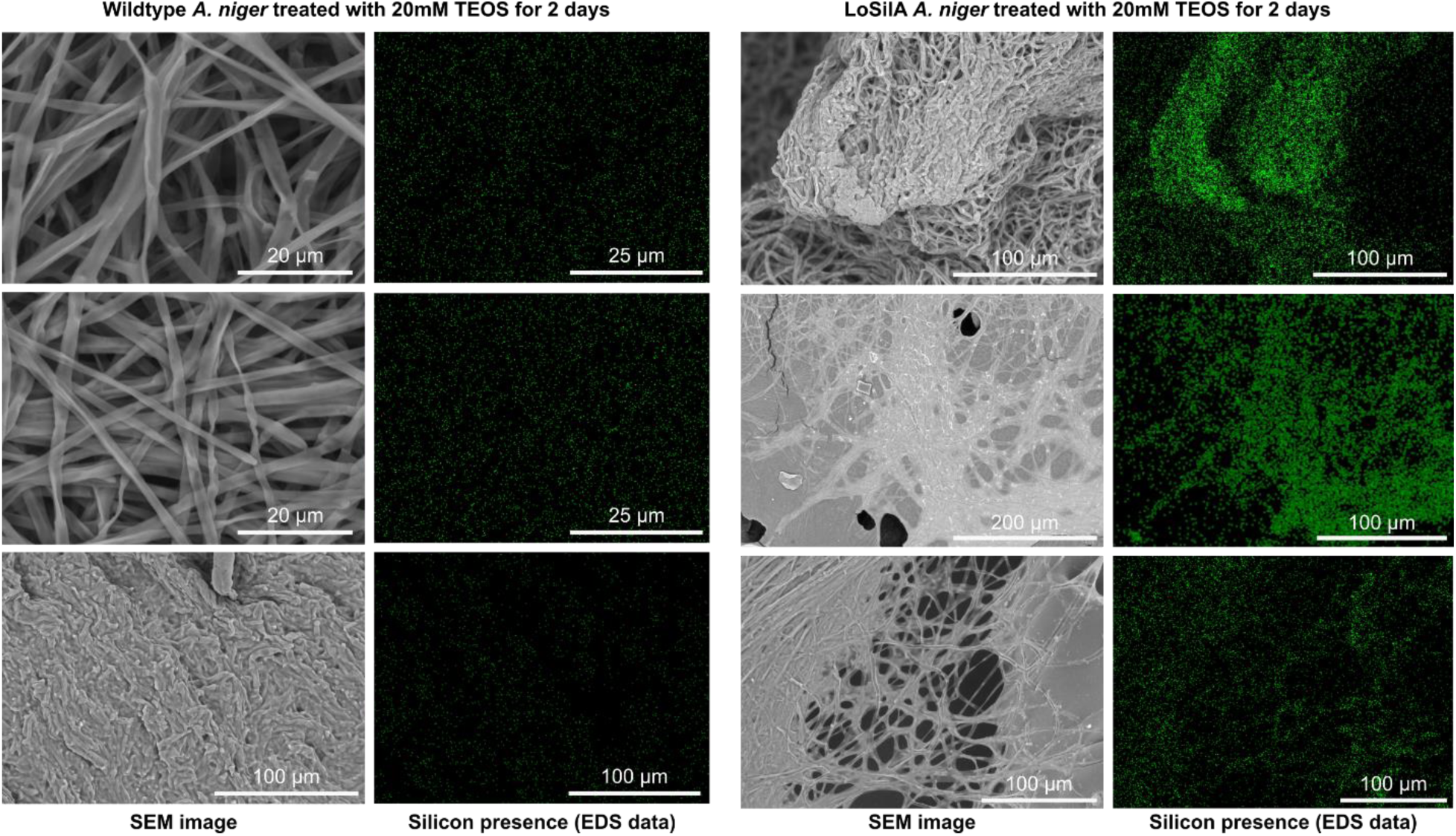
Representative SEM images with paired EDS data, where the green dots indicate the presence of a silicon signal, of wildtype (left) and LoSilA expressing fungi (right) treated with 20mM TEOS for 2 days showing polysilicate mineralization only on LoSilA expressing strains.

**Supplementary Figure S2.**
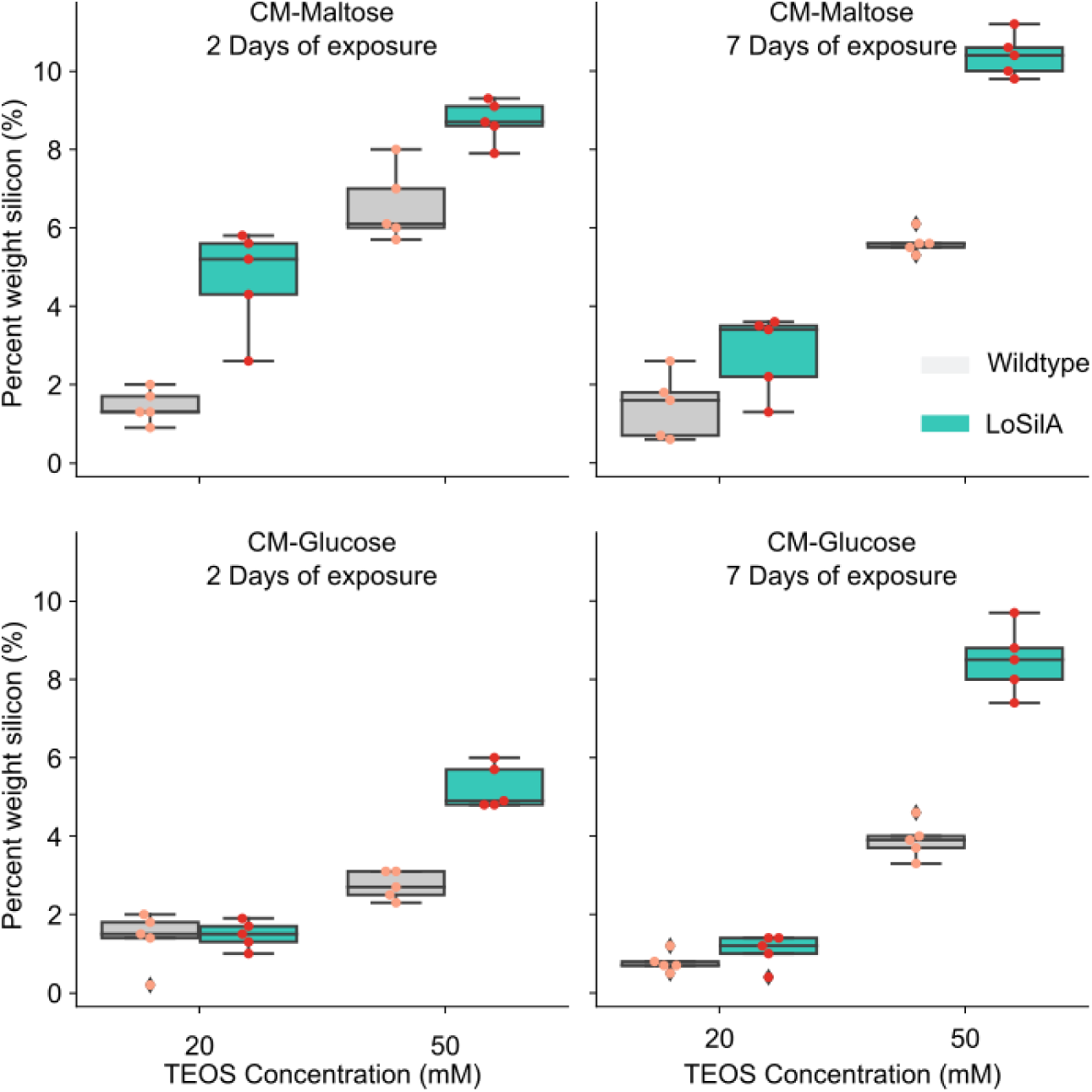
The degree of mycelial mineralization can be tuned. Boxplots summarizing EDS measurements of the percent weight of silicon on the surface of either the LoSilA expressing strain (teal) or a wildtype control (gray) grown in CM media supplemented with either maltose (top plots) or glucose (bottom plots) and then treated with either 20mM or 50mM TEOS for either 2 (left plots) or 7 (right plots) days.

**Supplementary Figure S3.**
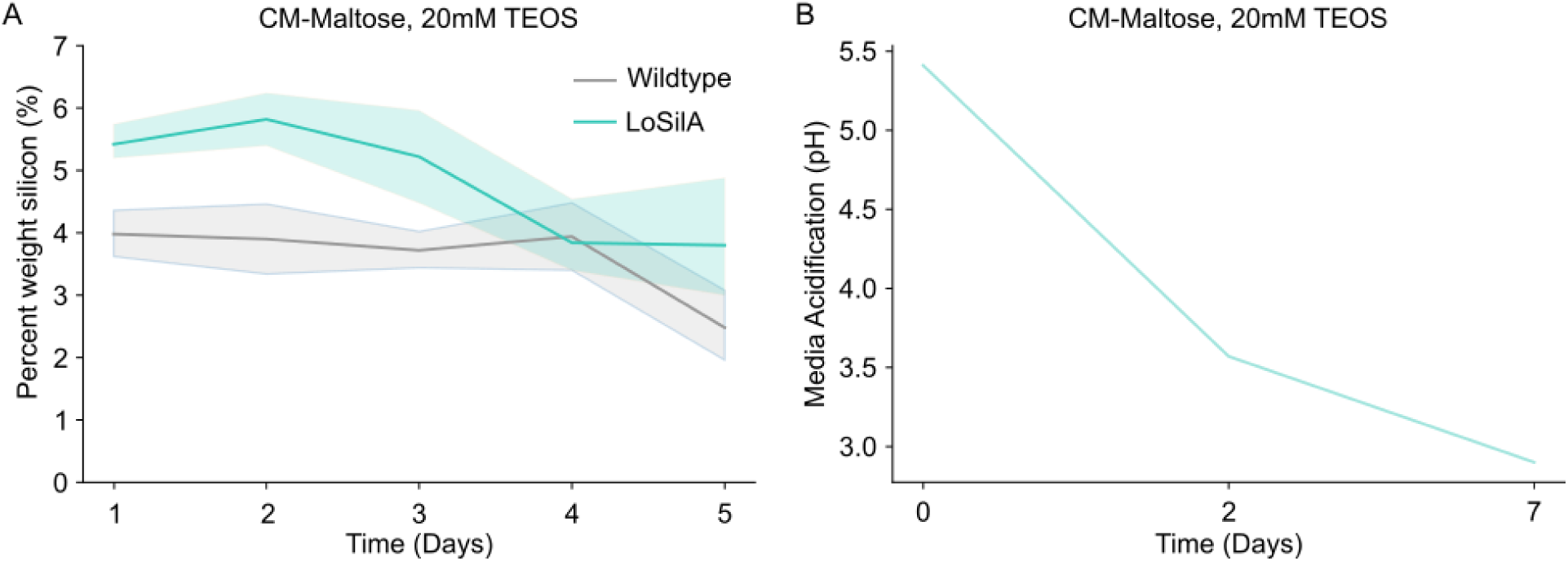
A) Line plot summarizing a daily time course of EDS measurements (n=5) of percent weight of silicon on the surface of LoSilA expressing (teal) or wildtype (grey) fungi. The shaded regions represent the 95% confidence interval. B) Line plot summarizing the pH of the media in the LoSilA time course cultures.

**Supplementary Figure S4.**
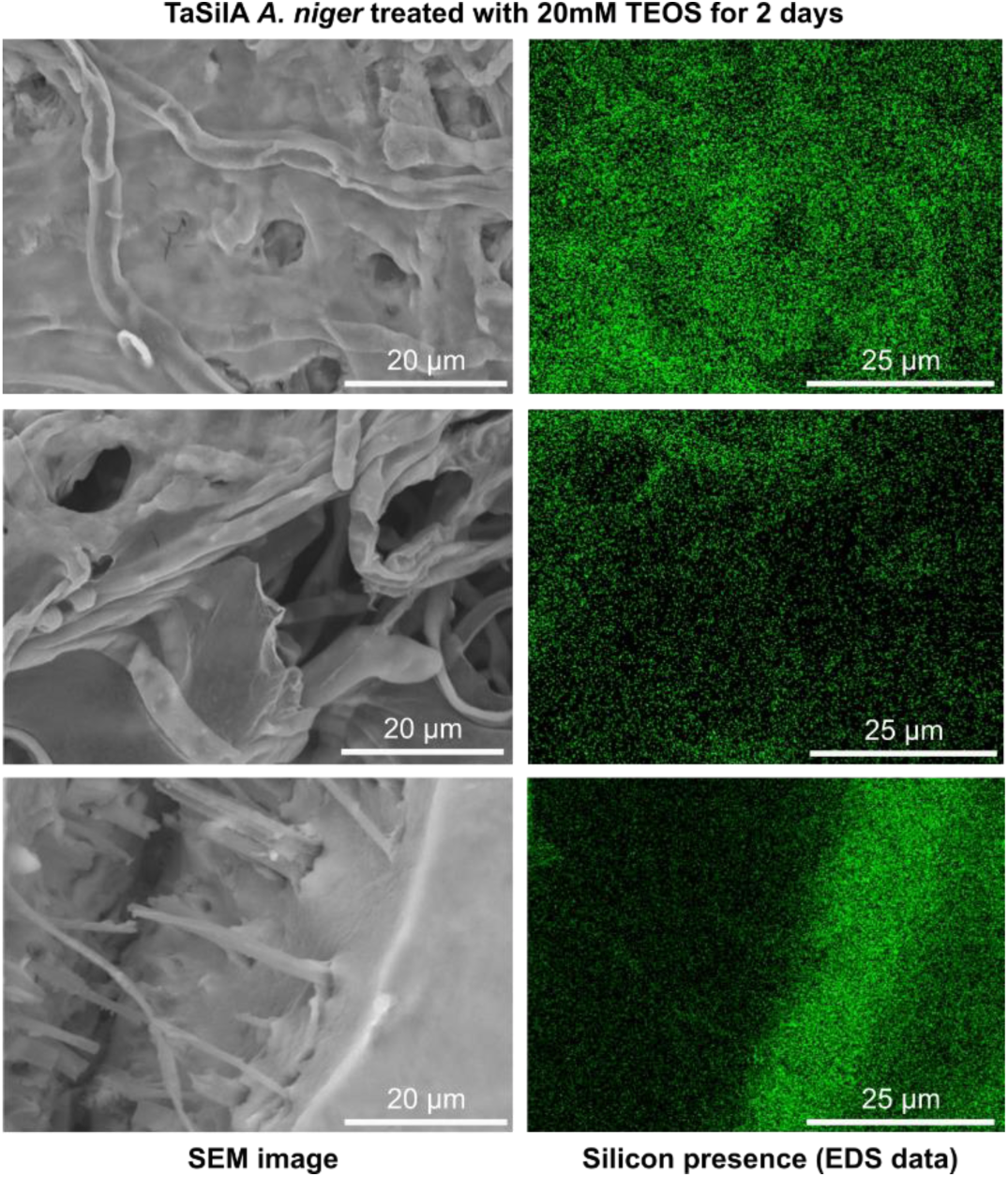
Representative SEM images with paired EDS data, where the green dots indicate the presence of a silicon signal, of TaSilA expressing fungi treated with 20mM TEOS for 2 days showing polysilicate mineralization.

**Supplementary Figure S5.**
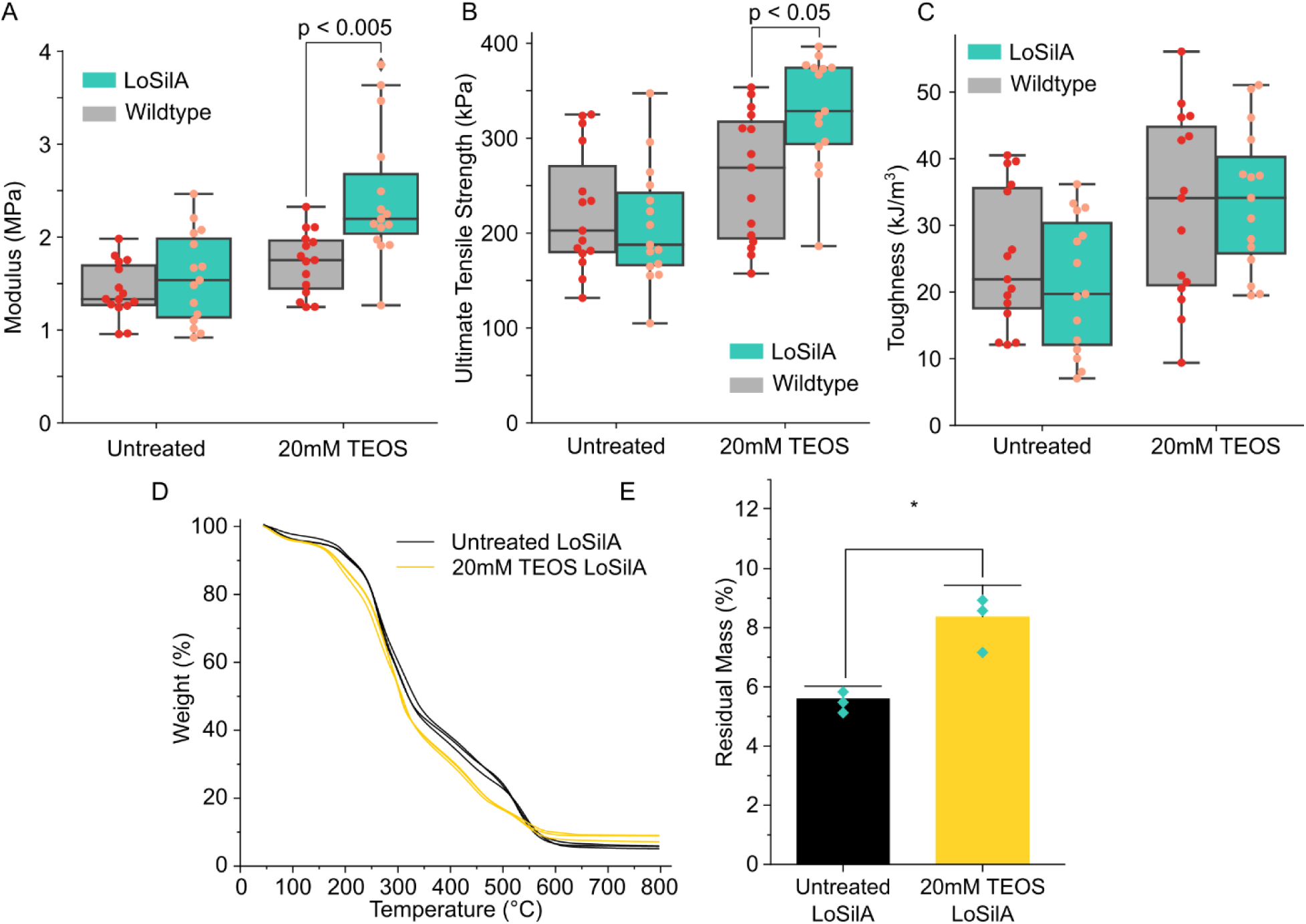
A-C) Box plots summarizing measurements of Young’s modulus, Ultimate tensile strength, and Toughness calculated from stress-strain curves of either a LoSilA expressing strain (teal) or a wildtype strain (gray) that is either treated with 20mM TEOS or that is untreated. Each dot represents an independent biological replicate. D) TGA curves showing Weight (%) vs Temperature of a LoSilA expressing strain that is either treated with 20mM TEOS (yellow) for 2 days or that is untreated (black). E) Bar plots summarizing residual mass (%) is taken from the TGA curves at the measurement endpoint (800⁰C).

**Supplementary Figure S6.**
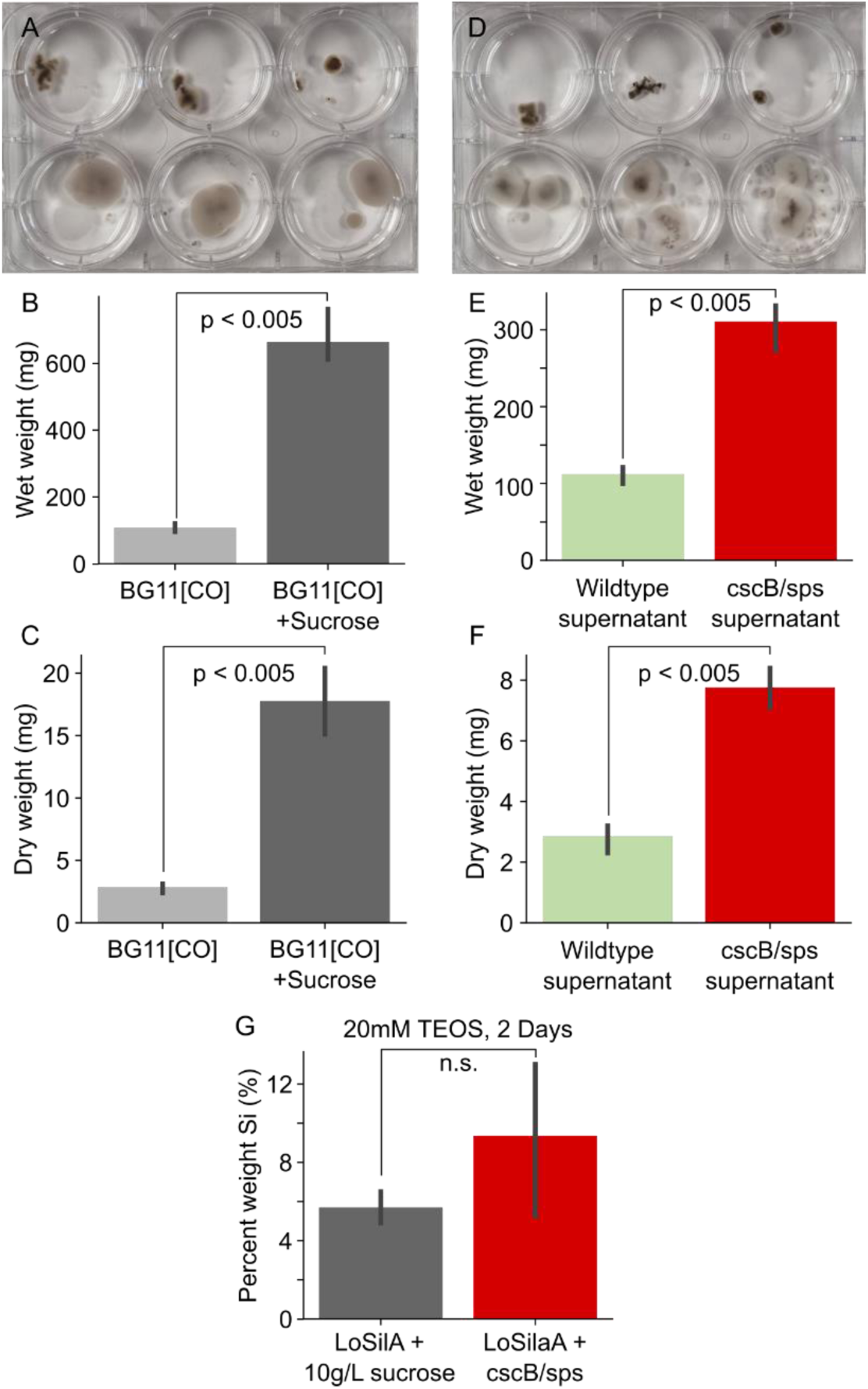
A) Representative image of LoSilA expressing *A. niger* grown in BG11[CO] media (top) or BG11[CO] supplemented with 10 g/L of sucrose (bottom). B,C) Bar plots summarizing measurements (n=3) of wet (B) and dry (C) weight of LoSilA expressing fungi grown in BG11[CO] media (light gray) or BG11[CO] supplemented with sucrose (dark gray). D) Representative image of LoSilA expressing fungi grown in the supernatant of wildtype *S. elongatus* cultures (top) or *cscB/sps S. elongatus* cultures (bottom). E,F) Bar plots summarizing measurements (n=3) of wet (B) and dry (C) weight of LoSilA expressing fungi grown in the supernatant of wildtype *S. elongatus* cultures (green) or *cscB/sps* expressing *S. elongatus* cultures (red). G) Bar plots summarizing EDS measurements (n=3) of the percent weight of silicon on the surface of LoSilA expressing fungi grown either using sucrose as a carbon source (gray) or in co-culture with *cscB/sps* expressing *S. elongatus* (red) and treated with 20mM TEOS for 2 days. “n.s.” indicates no statistically significant difference.

**Supplementary Figure S7.**
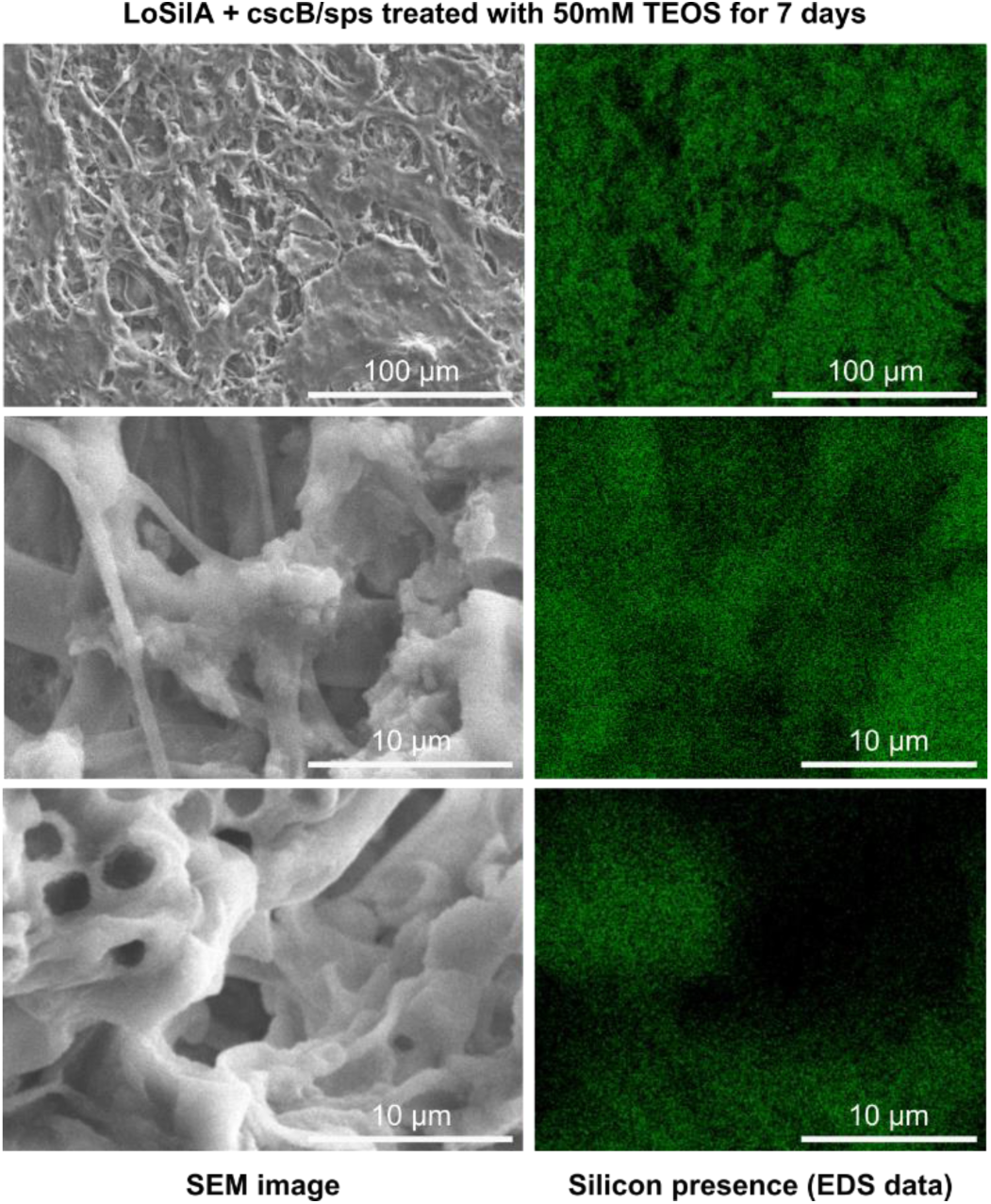
Representative SEM images with paired EDS data, where the green dots indicate the presence of a silicon signal, of LoSilA expressing fungi cocultured with cscb and sps expressing *S. elongatus* treated with 50mM TEOS for 7 days showing polysilicate mineralization.

